# FAM35A associates with REV7 and modulates DNA damage responses of normal and BRCA1-defective cells

**DOI:** 10.1101/287375

**Authors:** Junya Tomida, Kei-ichi Takata, Sarita Bhetawal, Maria D. Person, Hsueh-Ping Chao, Dean G. Tang, Richard D. Wood

## Abstract

In order to exploit the specific vulnerabilities of tumors, it is urgent to identify the basis of associated defects in genome maintenance. One unsolved problem is the mechanism of inhibition of processing of DNA double-strand break repair by REV7 and its influence on DNA repair pathways. We searched for REV7-associated proteins in human cells and found FAM35A, a protein of previously unknown function. By analyzing the FAM35A sequence we discovered that FAM35A has an unstructured N-terminal region and a C-terminal region harboring three OB-fold domains similar to single-stranded binding protein RPA. Knockdown of FAM35A caused sensitivity to DNA damaging agents, and FAM35A re-localized in damaged cell nuclei. In a BRCA1 mutant cell line, however, depletion of FAM35A increased resistance to camptothecin, suggesting that FAM35A participates in processing of DNA ends to allow more efficient DNA repair. Moreover, we found FAM35A absent in one widely used BRCA1-mutant cancer cell line (HCC1937) with anomalous resistance to PARP inhibitors. A survey of FAM35A alterations in cancer revealed that the gene is altered at the highest frequency in prostate cancers (up to 13%) and significantly less expressed in metastatic cases. The results reveal a new DNA repair factor with promise as a therapeutically relevant cancer marker.

---

REV7 is a multifunctional protein encoded by the *MAD2L2* gene in human cells. REV7 acts as an interaction module in several cellular pathways. One of its functions is as a component of DNA polymerase ζ, where it serves as bridge between the pol ζ catalytic subunit REV3L and the REV1 protein. A dimer of REV7 binds to two adjacent sites in REV3L by grasping a peptide of REV3L with a “safety-belt” loop^1,2^. REV7 protein is ~1000 times more abundant than REV3L^1^ and has additional functions and protein partners including chromatin-associated and post-translational modification proteins^3–8^). Complete ablation of REV7 gives rise to mice with defects in primary germ cells^6,9^. Recently studies uncovered a function of REV7 as a DNA resection inhibitor, limiting genomic repair by an unknown mechanism^10,11^. Although BRCA1 mutant cells are defective in homologous recombination, these studies found that one mode to partially restore recombination activity is by inactivation of REV7. It was proposed that REV7, together with 53BP1 and RIF1, inhibits 5’ DNA end resection to promote non-homologous end joining at the expense of homologous recombination.

To investigate novel functions and pathways involving REV7, we identified proteins associated with REV7 *in vivo*. We report here the first analysis of a previously uncharacterized REV7-interacting protein, FAM35A. We discovered that FAM35A is a novel factor that modulates the DNA damage sensitivity of normal and BRCA1-defective cells. Our analysis reveals that the C-terminal half of FAM35A contains three OB-fold domains similar to those in the single-stranded binding protein RPA large subunit. FAM35A has a disordered N-terminal portion, containing sites of DNA damage-dependent post-translational modification. Moreover, the *FAM35A* gene is deleted at an unusually high rate in prostate cancers, and in at least one well-studied BRCA1-defective breast cancer cell line. FAM35A is more weakly expressed in metastatic prostate cancers, suggesting it as an important marker for outcome and therapeutic decisions.

## Results and Discussion

### FAM35A interacts with REV7, 53BP1, and RIF1 *in vivo*

To isolate proteins associated with REV7, we engineered HeLa S3 cells that stably express REV7 with a C-terminal FLAG–HA epitope tag (REV7-FH). REV7-FH was sequentially immunoprecipitated from nuclear extract using FLAG and HA antibody beads^12,13^. This purified complex was separated by gradient gel electrophoresis and associated proteins from gel sections were identified by LC-MS/MS. We confirmed association with previously identified REV7 binding proteins including GLP ^6^, G9A ^6^, CAMP ^8^, GTF2I ^14^, POGZ ^7^, and HP1α ^7^ (Figure 1A and Table 1). A high-ranking, previously unstudied association was with the uncharacterized FAM35A protein (Figure 1A).

**Figure 1.**
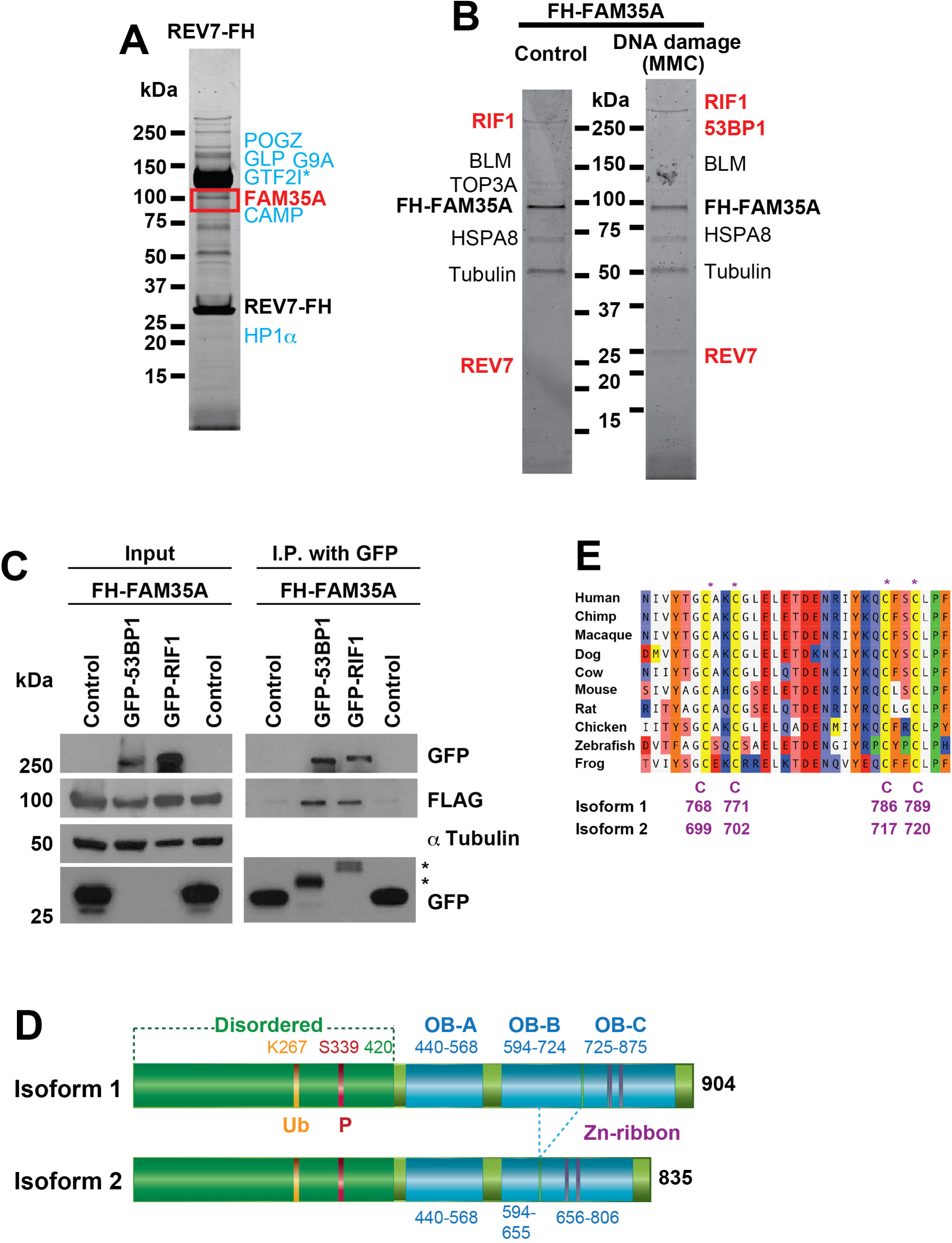
Identification of REV7 or FAM35A-associated proteins. Protein complexes were sequentially immunoprecipitated (using FLAG and HA antibody beads) from nuclear extracts of HeLa S3 cell lines stably expressing C-terminally FLAG–HA tagged REV7 (REV7-FH) or (N-terminally FLAG–HA tagged-FAM35A (FH-FAM35A). **(A)** REV7 complex and **(B)** FAM35A complex; associated proteins were identified by mass spectrometry. The FH-FAM35A complex was purified from HeLa S3 nuclear extracts after 18 h exposure to MMC (100 ng/mL). Proteins labeled in blue are previously published REV7 binding partners. Proteins labeled in red are involved in end-joining pathways of DSB repair. 4-20% gradient gels were stained with Sypro Ruby. (**C**) FH-FAM35A was co-transfected into human 293T cells with GFP empty vector (Control), GFP-53BP1 or GFP-RIF1. 48 h after transfection, cell lysates were made and used for immunoprecipitation with GFP antibody beads. After electrophoretic transfer of proteins, the membrane was cut into three sections to separate proteins >250 kDa (GFP as GFP-53BP1 and GFP-RIF1), 37-250 kDa (α-tubulin), and < 37 kDa (GFP as control) and immunoblotted with the indicated antibodies. Results for the input and immunoprecipitation (IP) product after gel electrophoresis are shown. The asterisk (*) in the IP lane marks degraded or truncated forms of 53BP1 and RIF1. (**D**) Domain schematic of human FAM35A derived from sequence prediction modeling. An N-terminal disordered region includes posttranslational modification sites. Locations of the three OB fold domains A, B, and C are shown, with a Zn-ribbon containing conserved Cys residues. One exon is absent in Isoform 2 compared to isoform 1, deleting 69 aa from OB domain B. (**E**) Multi-species alignment of a segment of FAM35A protein in the predicted Zn-ribbon. The 4 Zn-coordinating Cys residues (CxxC, CxxC), homologous to those in human RPA1, are evolutionarily conserved.

**Table 1.** DNA repair proteins and previously reported proteins identified in the REV7 complex by LC-MS/MS analysis

| Protein | Accession Number<br>(UniProtKB) | Molecular<br>Weight | Spectral<br>counts | Unique<br>peptides |
| --- | --- | --- | --- | --- |
| REV7 | Q9UI95\|MD2L2_HUMAN | 24 kDa | 453 | 15 |
| FAM35A | Q86V20-2\|FA35A_HUMAN | 92 kDa | 74 | 23 |
| GLP | Q9H9B1\|EHMT1_HUMAN | 141 kDa | 9 | 6 |
| G9A | Q96KQ7\|EHMT2_HUMAN | 135 kDa | 7 | 5 |
| CAMP | Q96JM3\|ZN828_HUMAN | 89 kDa | 127 | 29 |
| GTF2I | B4DH52\|B4DH52_HUMAN | 112 kDa | 1780 | 88 |
| POGZ | Q7Z3K3\|POGZ_HUMAN | 154 kDa | 147 | 31 |
| HP1α | P45973\|CBX5_HUMAN | 22 kDa | 8 | 3 |

To validate the association, a reciprocal experiment was performed by constructing a HeLa S3 cell line stably expressing FAM35A with an N-terminal FLAG–HA tag (FH-FAM35A). Cells were exposed to mitomycin C (MMC, 100 ng/mL) for 18 h, or mock-exposed. Following sequential immunoprecipitation with FLAG and HA antibody beads, proteins were separated and identified by mass spectrometry (Figure 1B). Proteins associating with FAM35A included REV7, RIF1, BLM and TOP3A (Table 2), with relatively more RIF1 peptides and 53BP1 identified following MMC exposure (Table 3).

**Table 2.** DNA repair proteins identified in the FAM35A complex by LC-MS/MS analysis

| Protein | Accession Number<br>(UniprotKB) | Molecular<br>Weight | Spectral<br>counts | Unique<br>peptides |
| --- | --- | --- | --- | --- |
| FAM35A | Q86V20-2\|FA35A_HUMAN | 92 kDa | 170 | 37 |
| REV7 | Q9UI95\|MD2L2_HUMAN | 24 kDa | 5 | 2 |
| RIF1 | Q5UIP0-2\|RIF1_HUMAN | 272 kDa | 6 | 4 |
| BLM | H0YNU5\|H0YNU5_HUMAN | 144 kDa | 6 | 4 |
| TOP3A | Q13472\|TOP3A_HUMAN | 112 kDa | 5 | 3 |

**Table 3.** DNA repair proteins identified in the FAM35A complex after MMC treatment by LC-MS/MS analysis

| Protein | Accession Number<br>(UniprotKB) | Molecular<br>Weight | Spectral<br>counts | Unique<br>peptides |
| --- | --- | --- | --- | --- |
| FAM35A | Q86V20-2\|FA35A_HUMAN | 92 kDa | 623 | 69 |
| RIF1 | Q5UIP0-2\|RIF1_HUMAN | 272 kDa | 315 | 112 |
| REV7 | Q9UI95\|MD2L2_HUMAN | 24 kDa | 21 | 4 |
| Ku80 | P13010\|XRCC5_HUMAN | 83 kDa | 6 | 4 |
| BLM | H0YNU5\|H0YNU5_HUMAN | 144 kDa | 6 | 4 |
| 53BP1 | Q12888-2\|TP53B_HUMAN | 214 kDa | 3 | 3 |
| Ku70 | P12956\|XRCC6_HUMAN | 70 kDa | 11 | 7 |

Although REV7 is known to cooperate functionally with 53BP1 to limit resection at DNA breaks, REV7 was not detected in 53BP1 immunocomplexes, and it is unknown how REV7 connects with 53BP1 *in vivo* ^11^. To verify an association of FAM35A with 53BP1 and the additional DNA end-resection control factor RIF1, FH-FAM35A was co-transfected with GFP-RIF1 or GFP-53BP1 and expressed in 293T cells. All proteins were expressed at the predicted molecular weights (Figure 1C, “input” lanes). Following immunoprecipitation with GFP antibody beads, FAM35A co-immunoprecipitated with recombinant RIF1 or 53BP1 (but not with the control vector, Figure 1C). These interactions suggest that FAM35A may functionally bridge 53BP1 and REV7 in human cells

### FAM35A is an OB-fold protein that changes localization following DNA damage

We found *FAM35A* orthologs are present in vertebrate genomes, but not in invertebrates or plants. Multiple protein isoforms arising from alternative splicing are annotated in genomics databases for human (Uniprot accession number Q86V20) and mouse FAM35A. Isoforms 1 and 2 are the most common, encoding 904 and 835 amino acid proteins, respectively. They arise by differential splicing of one in-frame exon (Fig. 1D). Both mRNA isoforms of FAM35A are ubiquitously expressed in different cell and tissue types (gtexportal.org).

BLAST searches for sequence homologs did not reveal significant primary sequence similarity to gene products other than FAM35A. We therefore analyzed the FAM35A protein sequence using PSI-BLAST structure prediction servers. The N-terminal half of the protein is predicted to be disordered up until about residue 420 (Figure 1D) and this region contains post-translational modification sites. Previous proteomic surveys identified a conserved ubiquitin modification^15^ and a conserved SQ site^16^ in FAM35A in which human S339 is phosphorylated after exposure to ionizing radiation or UV radiation^16^. With high significance, the C-terminal portion of FAM35A is predicted to contain three OB-fold domains structurally homologous to those in the 70 kDa subunit (RPA1) of the single-stranded DNA binding protein RPA (Figure 1D, Supplementary Figure 1). The three OB folds are similar to DNA binding domain folds A, B, and C of RPA^17,18^. OB fold domain C is predicted to include four conserved cysteine residues (Figure 1E) at the core of a zinc-binding ribbon, homologous to a loop in the same position in RPA (Supplementary Figure 1). Together, the OB folds and Zn ribbon form the elements of DNA binding and orientation enabling RPA1 to simultaneously bind to protein partners and to single-stranded DNA in an 8-10 nt binding mode^18–20^. The region including the 4-Cys Zn-ribbon identifies the domain of unknown function (PF15793) that is conserved in FAM35A homologs as annotated by the Pfam database. C-terminal helices are predicted present following OB domains A and B, in positions corresponding to helices involved in multimerization of the OB-folds in RPA subunits^17,18^ (Supplementary Figure 1).

We constructed a human U2OS cell line stably expressing GFP-FAM35A. Cells were exposed to MMC (100 ng/mL, 24 h) or mock–exposed, and then fixed and stained with DAPI and anti-GFP. In cells exposed to DNA damaging agent, GFP-FAM35A was concentrated into foci in the nucleus (Figure 2A, Supplementary Figure 4), suggesting a direct involvement in DNA repair.

**Figure 2.**
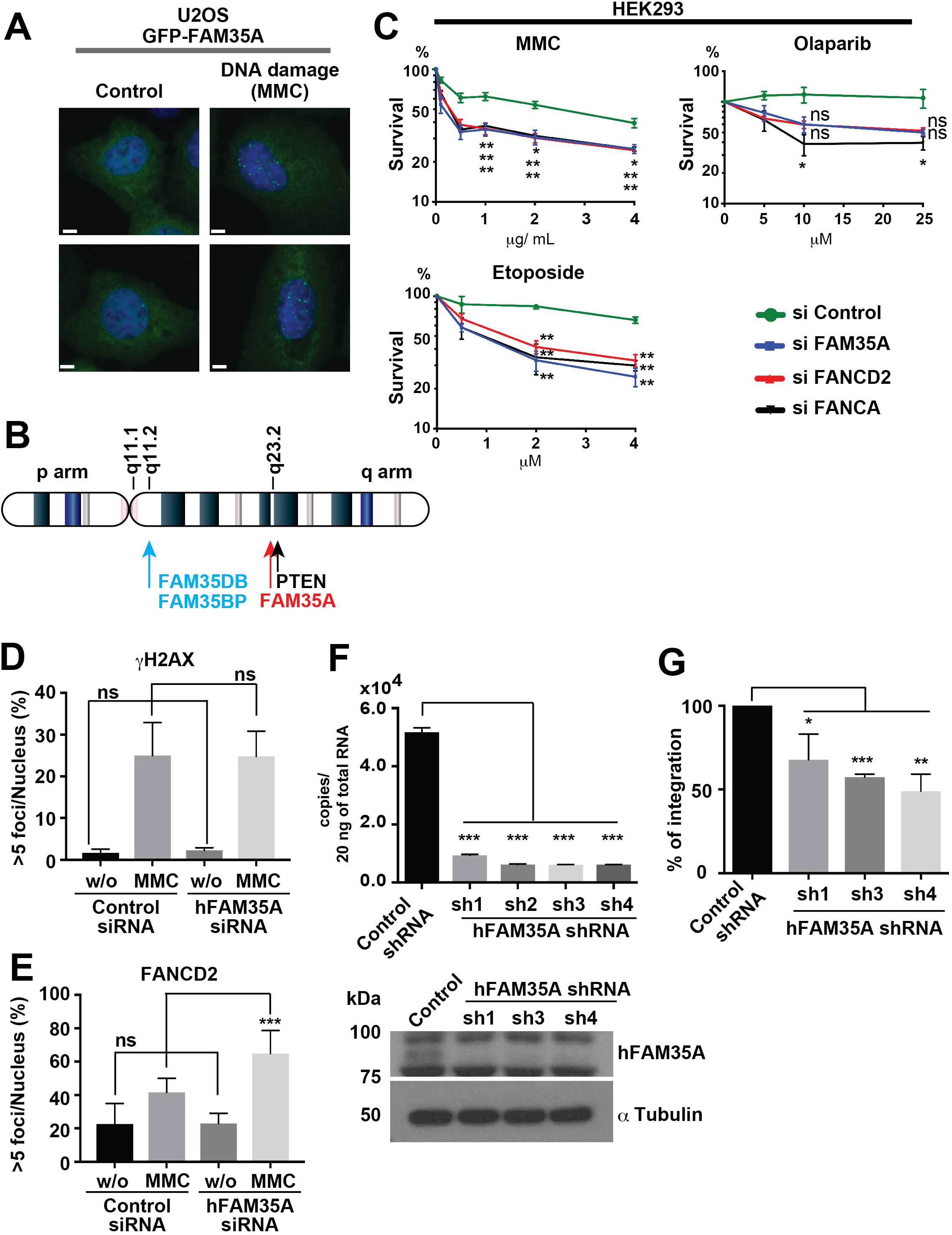
FAM35A is a DNA damage response gene. (**A**) GFP-FAM35A forms nuclear foci upon DNA damage. A U2OS cell line stably expressing GFP-FAM35A was exposed to MMC (100 ng/mL, 24 h) (right) or to mock treatment (left). The following day, cells were fixed and stained for DAPI and anti-GFP. Scale bars: 6 μm. (**B**) Two pseudogenes (*FAM35DP* and *FAM35BP*) with > 98% identity to *FAM35A* are located on chromosome 10q22. Both *FAM35DP* and *FAM35BP* are present in genomes of apes and old-world monkeys, but not in other mammalian genomes. By inference, these pseudogenes arose by whole gene duplication in the common ancestor of the Catarrhines about 25-30 million years ago. A third pseudogene (*FAM35CP*) is located on chromosome 14 (not shown). *FAM35CP* is an inactive spliced product of reverse transcription (> 95% identity) that was integrated into an intron of the galactosylceramidase (*GALC*) gene at 14q31.3. *FAM35CP* is present in apes but not old-world monkeys, indicating a more recent evolutionary origin. (**C**) Acute depletion of FAM35A causes hypersensitivity to several DNA damaging agents but not to olaparib. The survival of HEK293 cells, FAM35A acutely depleted and control, was monitored following exposure to MMC, etoposide, and olaparib. siControl (circle symbol, green line). siFAM35A (square symbol, blue line). siFANCD2 (triangle symbol, red line). siFANCDA (triangle symbol, black line). siRNA treated cells were plated and exposed to indicated dose of agent for 48 h. Cellular viability was measured 48 h later. Data represents mean ± SEM. n=3. (*) *P* < 0.05 and (**) P < 0.001 by unpaired *t*-test. (**D-E**) Quantification analysis of >5 γH2AX foci (**D**) and >5 FANCD2 foci (**E**) in FAM35A acutely depleted HEK293 cells. Cells with >5 foci/nucleus were counted after MMC (100 ng/mL, 24 h) or control treatment. (Data shown are means ± SE of more than 250 nuclei from two independent experiments. ns: not significant and (***) P < 0.0001 by unpaired *t*-test. (**F**) Depletion of endogenous FAM35A mRNA and protein from 293T cell lines carrying shRNA to FAM35A. Upper panel, mRNA levels. Lower panel, FAM35A protein expression. Cell lysates from stably FAM35A depleted 293T cell lines and control were made. After electrophoretic transfer of proteins, the membrane was immunoblotted with anti-FAM35A or anti-α tubulin. (**G**) Plasmid integration assay. pDsRed monomer plasmid with antibiotic resistance (Hygromycin B) was linearized by restriction enzyme. The linearized plasmid was transfected into stably FAM35A depleted 293T cell lines and control. Colonies were counted after antibiotic selection (Hygromycin B 300 μg/ml). Data represent mean ± SEM. n=4. (*) *P* < 0.05, (**) P < 0.001 and (***) P < 0.0001 by unpaired *t*-test.

### FAM35A-depletion sensitizes cell lines to DNA damage

The human *FAM35A* gene is located on chromosome 10q23.2. Three pseudogenes are also present in the human genome, two of them on 10q22 (Figure 2B) with high (> 98%) sequence identity to *FAM35A*. Precise nuclease-mediated knockout of human *FAM35A* is therefore challenging, as simultaneous targeting of pseudogenes would likely cause chromosome rearrangements and deletion. siRNA was used to acutely deplete FAM35A from human HEK293 cells and investigate its role in DNA repair. FAM35A-depleted HEK293 cultures were hypersensitive to MMC and etoposide, with sensitivity comparable to that conferred by depletion of Fanconi anemia *FANCA* and *FANCD2* gene expression (Figure 2C). In HEK293 cells, *FAM35A*-siRNA did not sensitize to the PARP inhibitor olaparib (Figure 2C).

We investigated the impact of acute depletion of FAM35A on other markers of DNA damage responses in HEK293 cells. Cells with >5 γH2AX foci or >5 FANCD2 foci were quantified after MMC exposure (100 ng/mL, 24 h) and in control cells. MMC exposure induced cellular γH2AX foci as expected; there was no significant change in this pattern in cells depleted for FAM35A, indicating intact signaling leading to γH2AX formation (Figure 2D). The Fanconi anemia signaling pathway was also intact in FAM35A-depleted cells, showing about 1.5-fold more foci after MMC exposure compared to cells with siControl (Figure 2E).

Depletion of REV7 reduces the efficiency of non-homologous end joining (NHEJ) ^10,11^ by promoting resection that channels repair of DNA double-strand breaks into homologous recombination and other pathways. Because of the association between FAM35A and the resection control factors REV7, RIF1 and 53BP1 (Figure 1A-C), we investigated whether FAM35A depletion affects end joining, using a plasmid integration assay ^10^. REV7, 53BP1 and RIF1 depletion decreased integration ratio in this assay. We confirmed efficient knockdown of *FAM35A* mRNA and protein in 293T cells using qPCR and immunoblot analysis (Figure 2F). The plasmid integration ratio decreased significantly after FAM35A depletion (Figure 2G), suggesting that FAM35A is involved in modulating double-strand break repair pathway choice.

### FAM35A deficiency in BRCA1-deficient cells and resistance to camptothecin and PARP inhibitors

In BRCA1 mutant cells, REV7 depletion restores homologous recombination (HR) by restoring 5’ end resection^10,11^. We hypothesized that FAM35A depletion from BRCA1 mutant would increase resistance to a DNA strand-breaking agent. We engineered a BRCA1 mutant cell line (MDA-MB-436) expressing FAM35A shRNA. Efficient knockdown was verified by qPCR (Figure 3A). The FAM35A-depleted BRCA1 mutant cells and controls were assayed for sensitivity to camptothecin using a colony forming assay. FAM35A-depletion from the BRCA1 mutant cell line significantly alleviated the sensitivity to camptothecin (Figure 3A). We investigated a further widely used cell line, HCC1937, which has a known inactivating mutation in BRCA1 (Figure 3B). Multiple studies have noted that HCC1937 is anomalously resistant to PARP inhibitors in comparison to other BRCA1 mutant cell lines, for unknown reasons^21–23^. In HCC1937, RAD51 foci (a marker of DNA end resection) are generated after DNA damage exposure ^24,25^, and homologous recombination is partially active despite BRCA1mutation^25^. We found that *FAM35A* mRNA expression is absent in HCC1937 cells (Figure 3C). FAM35A expression was present in HCC1937BL, an EBV-immortalized lymphoblastoid cell line from the same patient (Figure 3C). HCC1937BL is heterozygous for the BRCA1 mutation, indicating loss of heterozygosity for both FAM35A and BRCA1 in the HCC1937 mammary cancer patient.

**Figure 3.**
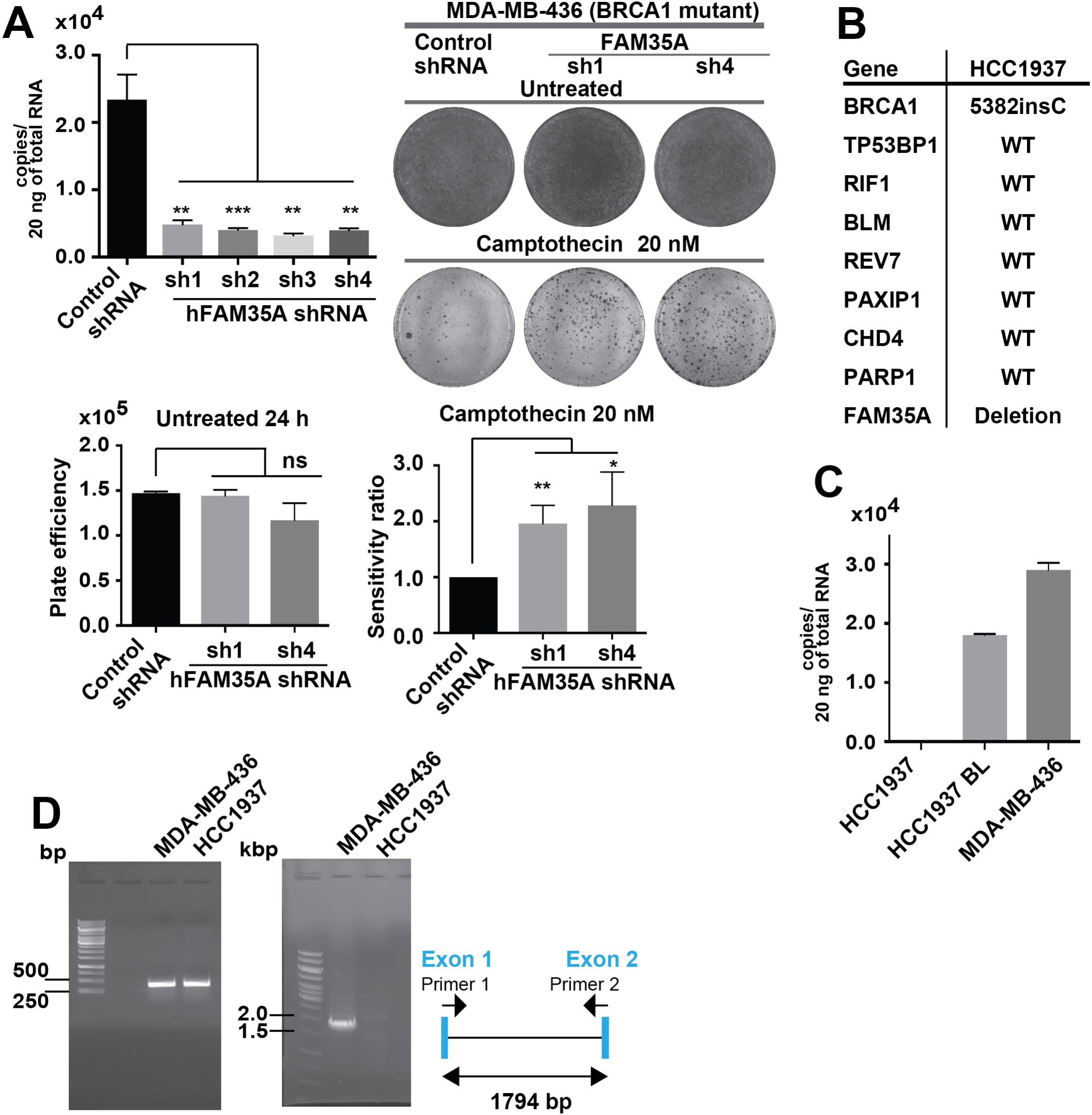
Increased resistance to camptothecin following FAM35A depletion / KO in BRCA1-deficient cells. (**A**) Quantification of depletion of endogenous FAM35A mRNA and protein from 293T cell lines carrying shRNA to FAM35A. Colony forming assay: MDA-MB-436 cells infected with non-targeting control or *FAM35A* shRNAs were treated for 24 h with 20 nM camptothecin, with medium changed and incubated until colonies appear; cells were then fixed and stained. Top row is untreated; second row is treated with camptothecin. Colony and cell numbers were counted. Bottom left is the number of cells 24 h after seeding 1.5 ×10^5^ cells. Bottom right is the colony count. Data represent mean ± SEM. n=3. ns: not significant, (*) *P* < 0.05 and (**) P < 0.001 by unpaired *t*-test. (**B**) Genes related to PARP inhibitor resistance in BRCA1 mutant cells, and alterations of these genes in the HCC1937 cell line according to the CCLE. (**C**) Quantification of endogenous FAM35A mRNA expression. Endogenous FAM35A mRNA from HCC1937 (*BRCA1* mutant), HCC1937BL and MDA-MB-436 (*BRCA1* mutant) cell lines was quantified using qPCR. Data represent mean ± SEM. n=3. nd: not detectable and (****) P < 0.0001 by unpaired *t*-test. (**D**) Detection of FAM35A Exon 1 & 2 deletion in genomic DNA using PCR. Deletion of FAM35A exon 1 & 2 is confirmed in the HCC1937 cell line. In HCC1937 BL, endogenous FAM35A mRNA expression level was detectable. PCR primers for amplification of genomic DNA were designed in exon 1 (forward; primer 1) and exon 2 (reverse; primer 2). The predicted PCR product size is 1794 bp.

Genomic analysis of DNA from HCC1937 and MDA-MB-436 cells showed that *FAM35A* DNA flanking exons 1 and 2 was not present in HCC1937 cells (Figure 3D). This is consistent with a deep deletion at the *FAM35A* locus, notated by the Cancer Cell Line Encyclopedia (CCLE, Broad Institute) in HCC1937 cells (Supplementary Figure S2). Reported genes related to PARP inhibitor resistance in BRCA1 mutant cells include *53BP1*, *MAD2L2*/*REV7*, *RIF1*, *BLM*, *CHD4* and *PTIP* ^10,11^ ^26,27^. None are deleted in HCC1937 (Figure 3B). DNA double-strand break repair factors ATM ^10^, NBS1 ^10^, and RNF8 ^10^ are also preserved in HCC1937 according to the CCLE.

These data suggest an involvement of FAM35A in resection inhibition in parallel with REV7, 53BP1 and RIF1 ^10,11^. Like 53BP1-depleted BRCA1 mutants ^26,28^, FAM35A-depleted BRCA1 mutant cells acquire camptothecin resistance. As HCC1937 cells can form 53BP1 foci and Mre11 foci^25^ following DNA damage ^24^, FAM35A appears to operate downstream of these factors and may serve as a link between 53BP1 and REV7. It is notable that the HCC1937 cell line relies on pol θ-dependent alternative end-joining for survival, perhaps unusually so for a BRCA1-defective cell line ^29^. A likely explanation is that the excessive DNA end resection activity arising from the FAM35A defect channels a substantial fraction of breaks into repair by DNA polymerase θ-mediated alternative end-joining ^30^.

### FAM35A alterations and implications for cancers

Currently, there are two major models to account for PARP inhibitor resistance in BRCA1 mutant cells: 1) 5’ resection through dysfunction of 53BP1, RIF1 or REV7 ^10,11^ and 2) deletion of recruiters (PTIP, CHD4 and PARP1) of enzymes that degrade newly synthesized DNA^31^. Our discovery of FAM35A deletion in HCC1937 alongside reported evidence suggests a plausible mechanism of FAM35A-associated PARP inhibitor resistance in line with the first model, via 5’ resection.

We investigated the incidence of FAM35A alterations in cancer using cancer genomics databases. Mutations and copy number changes are found in FAM35A at a relatively low level (< 5% of cases) in most cancer types. Strikingly, however, prostate cancer (PCa) data show deletion of *FAM35A* in a high fraction (6-13%) of cases (Figure 4A). Prostate cancers are the only cancer type with this level of FAM35A deletion (for cancers with at least 40 cases per dataset). The *PTEN* gene, frequently deleted in PCa, is located on 10q23.2 about 0.5 Mb distal of *FAM35A* (Figure 2B). Some cases of FAM35A deletion are independent of *PTEN* deletion (Figure 4B). A meta-analysis of 6 datasets containing *FAM35A* mRNA expression data was performed to compare metastatic and primary tumors in prostate cancer. *FAM35A* mRNA expression is significantly reduced in metastatic prostate cancer (Figure 4C). In prostate cancers, deletion of FAM35A is essentially exclusive from mutation or deletion of BRCA1. From the data presented here, tumors with FAM35A deletion might be predicted to be susceptible to treatment with DNA crosslinking or double-strand break inducing agents, but not to olaparib. Finally, we note that some other commonly used cancer cell lines harbor genetic deletion for FAM35A as annotated in the CCLE, although each must be confirmed experimentally. We found that prostate cancer cell line LNCAP clone FGC still expresses FAM35A mRNA (Supplementary Figure 2).

**Figure 4.**
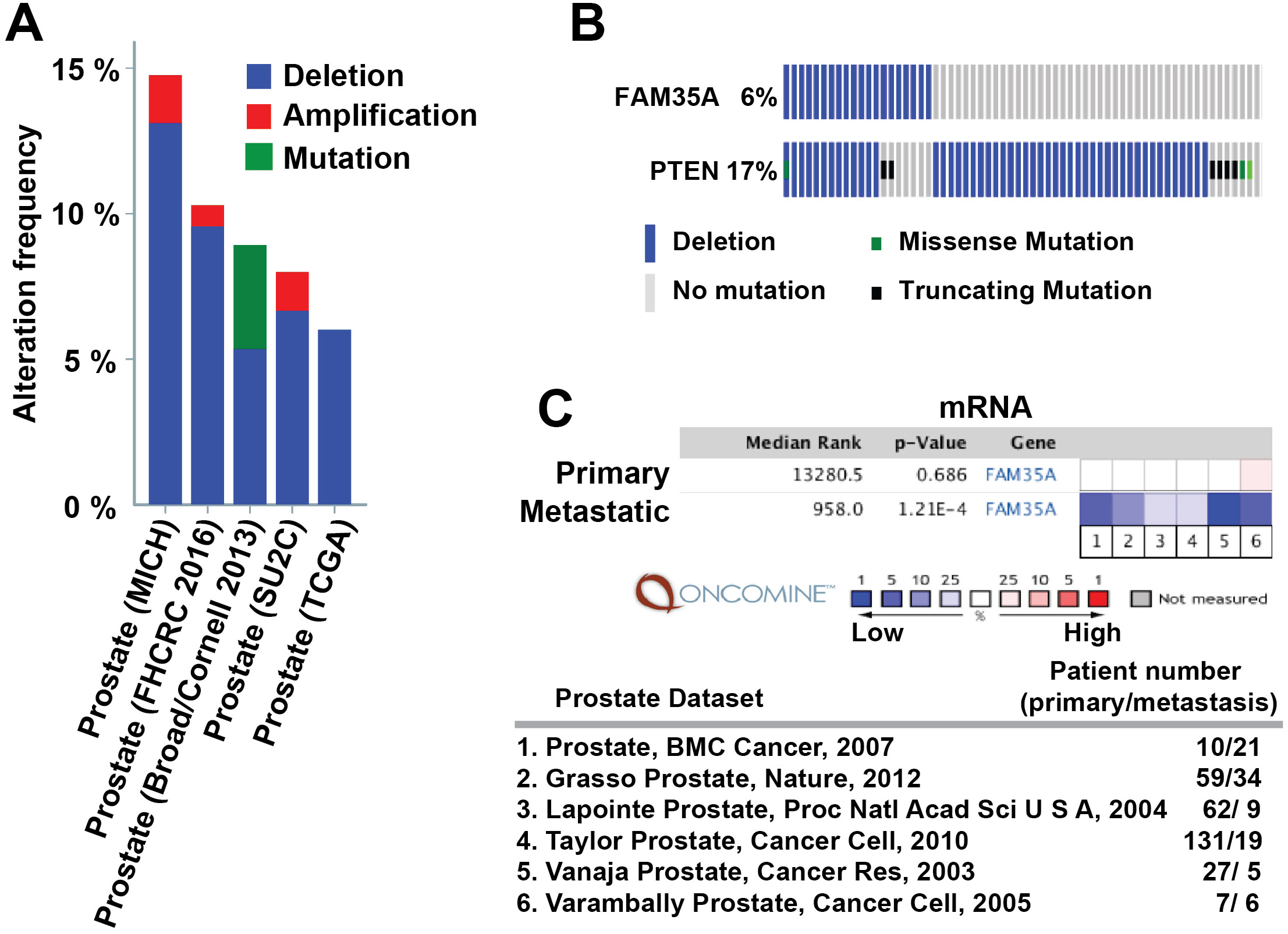
FAM35A is a candidate tumor/metastasis suppressor gene in prostate cancer. (**A**) Prevalence of *FAM35A* deletion in prostate cancer (PCa) genomes analyzed via the cBioPortal. FAM35A gene is depleted in ~6-13% of PCa, in the studies indicated. (**B**) FAM35A deletion is sometimes independent of alteration in PTEN. Comparison of *FAM35A* and *PTEN* alterations in PCa according to TCGA data (333 samples, Cell, 2015) from the cBioPortal. Blue bar, homozygous deletion. Gray bar, no mutation. Green square, missense mutation. Black square, truncating mutation. (**C**) *FAM35A* mRNA expression in primary vs. metastatic tumor according to Oncomine. The *p*-value for a gene is its *p*-value for the median-ranked analysis. The enumerated studies are Prostate BMC Cancer ^45^, Grasso Prostate ^46^, Lapointe Prostate ^47^, Taylor Prostate ^48^, Vanaja Prostate ^49^, Varambally Prostate ^50^.

The data presented here strongly implicate FAM35A in inhibition of DNA end resection following strand breakage (Supplementary Figure 3). In BRCA1 mutant cells, knockout of FAM35A allows recombination and/or alternative end-joining repair pathways to be utilized more efficiently, and helps explain the relative resistance to PARP inhibitors. Based on the protein interactions uncovered here and our prediction of single-stranded DNA binding activity, several possible mechanisms for FAM35A can be explored. The protein may interact directly with single stranded DNA via its OB fold domains and block activity of resection enzymes (BLM-DNA2, BLM-EXO1, and Mre11); alternatively, FAM35A could associate with BLM to prevent the interaction of BLM with DNA2 or EXO1.

## METHODS

### Human cell cultures and transfections

HEK293 (ATCC CRL-1573), U2OS (ATCC HTB-96) and HEK293T (ATCC CRL11268) cells were maintained in Dulbecco’s Modified Eagle’s medium GlutaMAX™ (Life Technologies, Carlsbad, CA) supplemented with 10% fetal bovine serum and penicillin/streptomycin in a 5% CO_2_ incubator at 37 °C. Human HeLa S3 cells were maintained in RPMI supplemented with 10% fetal bovine serum and penicillin/streptomycin in a 5% CO_2_ incubator at 37 °C. All cell lines were routinely checked for mycoplasma contamination using the MycoAlert detection kit (Lonza). Cell lines were validated by STR DNA fingerprinting by the Cell Line Identification Core of the MD Anderson Cancer Center. The STR profiles were compared to known ATCC fingerprints (ATCC.org), to the Cell Line Integrated Molecular Authentication database (CLIMA) version 0.1.200808 (http://bioinformatics.istge.it/clima/)^32^, and to the MD Anderson cell line fingerprint database.

### siRNA transfection

siRNA transfections were performed with Lipofectamine RNAiMAX (Life Technologies) according to manufacturer instructions and as described previously ^33,34^. The messenger RNA target sequences used for siRNAs were as follows: FANCA-specific RNAs (5’-AAGGGUCAAGAGGGAAAAAUA-3’)(Invitrogen), FANCD2-specific RNAs (5’-AAUGAACGCUCUUUAGCAGACAUGG-3’)(Invitrogen), FAM35A-specific RNA (5’-AAGGAGUGGUUCUGAUUAA-3’) (Thermo Scientific), #6 (5’-GUACUAAGAGUUGUUGAUU -3’) (Thermo Scientific), #7 (5’-GAACAGGAUCUACAAACAA -3’) (Thermo Scientific) and #8 (5’-GCUCACAGUUCUCUGAAGA -3’) (Thermo Scientific). ON-TAR-GETplus Non-Targeting siRNAs were used as a negative control designated “siControl” (Thermo Scientific). The siRNAs were introduced into HEK293 cells.

### Immunoprecipitation assay and immunoblotting

Immunoprecipitation was performed as described ^33,34^. 48 h after transfection (for FH-FAM35A and GFP, GFP-53BP1 or GFP-RIF1) in 293T cells, the cells were harvested. Each cell pellet was suspended with 300 µL of 0.5B (500 mM KCl, 20 mM Tris-HCl [pH 8.0], 5 mM MgCl_2_, 10% glycerol, 1 mM PMSF, 0.1% Tween 20, 10 mM β-mercaptoethanol), frozen in liquid nitrogen, thawed on ice, and sonicated. After centrifugation, 900 µL of 2B (40 mM Tris-HCl [pH 8.0], 20% glycerol, 0.4 mM EDTA, 0.2% Tween 20) was added to the supernatant and incubated with 10 µL of GFP agarose (MBL) for 4 h at 4°C. The bound proteins were washed with 700 µL of 0.1B (100 mM KCl, 20 mM Tris-HCl [pH 8.0], 5 mM MgCl_2_, 10% glycerol, 1 mM PMSF, 0.1% Tween 20, 10 mM β-mercaptoethanol), three times and eluted with 30 µL of 2x SDS loading buffer (100 mM Tris-HCl [pH 6.8], 4% SDS, 0.2% bromophenol blue, 20% glycerol, 200 mM DTT). These samples were separated by polyacrylamide gel electrophoresis, transferred to a membrane, and detected with the indicated antibodies and ECL reagents (GE Healthcare).

### Random plasmid integration assay

Plasmid integration assay was performed as previously described ^10,35,36^. Plasmid transfections were performed with Polyethylenimine 25kD linear from Polysciences (cat# 23966-2) as described^37^. Briefly, 24 h after transfection (with linearized pDsRed-Monomer-Hyg-C1 (Clontech; #632495) with SmaI or XhoI) in FAM35A-depleted 293T cell lines and control cells, 1.5 ×10^5^ cells were plated into replicate 10 cm dishes for Hygromycin B treatment and 5 × 10^2^ cells were plated into replicate 10 cm dishes for plating efficiencies. The next day, Hygromycin B 300 μg/ml was added to the dish, and the cells were incubated until appearance of colonies (10 to 14 days). Cells were then fixed, stained, and counted. Statistical analysis performed by unpaired t-test (p < 0.05).

### shRNA vectors

shRNA vectors were purchased from MD Anderson core facility: shFAM35A#1 (V2LH2S_221094), shFAM35A#2 (V2LH2S_277883), shFAM35A#3 (V3LH2S_359959), shFAM35A#4 (V3LH2S_359963) and shScramble (RHS4346).

### Antibodies

Antibodies purchased from Sigma-Aldrich, with dilutions for immunoblotting, were: F3165, monoclonal anti-FLAG 1:10,000; F7425, monoclonal anti-aTubulin 1:8,000; T5168, HRP (horseradish peroxidase) conjugated anti-mouse IgG 1:10,000; A0168, HRP conjugated anti-rabbit IgG 1:10,000; HPA036582, polyclonal anti-FAM35A 1:200. D153-8 agarose conjugated anti-GFP (RQ2) was purchased from MBL International Corporation. From Clontech, polyclonal anti-GFP antibody 632592 was used at 1:200 (for immunostaining); monoclonal anti-GFP 632381, 1:3000.

### Mass Spectrometry

Protein identification was performed by mass spectrometry as described previously ^38^.

### DNA constructs

Human *FAM35A* cDNA (Clone ID: 6146593) was obtained from Thermo Scientific. Isoform 2 cDNA was PCR amplified from FAM35A cDNA as a *Xho*I-*Not*I fragment with 5’FAM35A (*Xho*I) primer (5’-CCGCTCGAGATGAGTGGAGGATCTCAAGTCCAC) and 3’FAM35A (*Not*I) primer (5’-TAAAAGCGGCCGCTCAGAGACGGGCATT-GGCTCCATGC) to clone into pETDuet-1 and pRSFDuet-1 (Novagen). Full-length REV7 cDNA was cloned into pETDuet-1 ^1^. The *Xho*I-*Not*I fragments from FAM35A and REV7 were inserted into pOZN and pOZC ^39^ (kindly provided by Hank Heng Qi, Children’s Hospital, Boston). After construction, expression vectors were confirmed by DNA sequencing. pDESTpcDNA5-FRT/TO-eGFP-Rif1 and pcDNA5-FRT/TO-eGFP-53BP1 were gifts from Daniel Durocher (Addgene plasmid # 52506 and # 60813 respectively) ^40,41^.

### DNA damage sensitivity in FAM35A-depleted cell lines

To analyze sensitivity to chemical DNA damaging agents, HEK293 were plated into white 96-well plates (1,250 cells/well). The next day, various concentrations of MMC, etoposide and olaparib were added to the wells, and the cells were incubated for 48 h. The cells were lysed, a reagent that luminesces in the presence of ATP was added (ATPLite One Step, Perkin Elmer). Luminescence was measured using a plate reader (Biotek Synergy II) and normalized to undamaged control. For the clonogenic assay, 1.5 × 10^5^ cells were plated in 10 mm culture plates and incubated for 24 h prior to DNA damage induction. Groups of plates were exposed to camptothecin (20 nM). After 24 h, medium was changed and cells were incubated until appearance of colonies (14 to 21 days). Cells were then fixed, stained, and counted. Statistical analysis performed by unpaired t-test (p < 0.05).

### Foci analysis in U2OS cells stably expressing GFP-FAM35A and FAM35A-depleted cell lines

Immunofluorescence photos were taken on a Leica DMI6000B microscope as described previously ^38^. The cells were plated into an 8-well chamber slide; the following day, cells were fixed with paraformaldehyde after MMC treatment (100 ng/ml 24 h) and stained for DAPI and GFP ^33^. To measure the formation of DNA double-strand breaks and downstream signaling, FAM35A acutely depleted and control HEK293 cells were plated in 8-well chamber slides, treated with MMC (100 ng/ml 24 h), fixed with paraformaldehyde, and stained for DAPI and γH2AX or FANCD2. γH2AX or FANCD2 foci numbers were counted and plotted.

### Genomic PCR

Total DNA was isolated from HCC1937 and MDA-MB-436 cells using DNeasy Blood & Tissue Kit (Qiagen) following manufacturer’s protocol. Total DNA (400 ng) was used as template for genomic DNA amplification of FAM35A exon 1 and 2 by PCR with Primer 1 (5’-ACGGGCGGCCGGATTTGCCCGGAGG) and Primer 2 (5’-CTTAATCTTTTGACCATCAAGG). For control, primers were REV3L-F (5’-ACAGCTTCAGAGGAAAGCCA) and REV3L-R (5’-GTTACCGATCGTGTCCGTTT).

### Quantitative PCR (qPCR) assay

Total RNA was extracted using RNeasy Mini Kit (Qiagen) following manufacturer’s protocol including on-column DNase treatment. Total RNA (1 µg) was used as template to synthesize cDNA with the High-Capacity cDNA Reverse Transcription Kit (Thermo Fisher). qPCR was then performed on the ABI 7900HT Fast Real-Time PCR System (Applied Biosystems). TaqMan primers and the probe sets for *hFAM35A* were purchased from Thermo Fisher (assay ID Hs04189036_m1). The absolute quantity (AQ) of transcripts for *hFAM35A* was determined using the generated standard curves. Standard curves for each gene were determined using the plasmid pRSFDuet-1 carrying one of *hFAM35A*. Each reaction was performed in triplicate.

### Bioinformatics

Analysis of predicted protein disorder used the DisEMBL (http://dis.embl.de), Pondr (http://pondr.com) and Phyre2 web-based servers ^42^. Protein structure prediction was cross-checked using multiple tools. Several structure prediction servers identified models corresponding to OB domain A. The ModWeb server for (https://modbase.compbio.ucsf.edu) predicts high confidence similarity (sequence identify 18%) of FAM35A residues corresponding to the OBA fold with PDB code 2K50 (*Methanobacterium thermoautotrophicum* RPA protein OB fold) and other with other RPA models. SwissModel (https://swissmodel.expasy.org/) located homology of FAM35A 904 aa isoform residues 583-735 (OB fold B) with human RPA1 OB-fold PDB code 1FGU. When the C-terminal 480 amino acids of FAM35A were submitted to Phyre2 for structure prediction, the highest scoring template was PDB code 4GOP (71% coverage, 99.4% confidence). OB folds A, B, and the first part of OB folds C were modeled from 4GOP. The OB fold C model and is consistent with a partial OB fold predicted for FAM35A residues 736-836, PDB code 3U50.

Cancer genome data and Cancer Cell Line Encyclopedia data was accessed from the cBioPortal for Cancer Genomics ^43^. Oncomine^44^ (www.oncomine.com) was used for meta-analysis of 6 datasets containing FAM35A mRNA expression data (mostly microarrays) with comparisons between metastatic and primary tumor in prostate cancer. Total patient numbers and detailed information regarding published datasets and associated publications are indicated in Figure 4B.

## Author Contributions

JT and KT conceived and designed the experiments. JT and RDW designed the research. JT and SB, MDP performed research. JT, KT, H-PC, DG, and RDW analyzed data. JT and RDW wrote the paper.

## Acknowledgements

We thank Edmund O’Brien, Sara K. Martin, Megan Lowery, Karen Boulware for assistance with experiments, Mary Walker for editorial help, Dr. Stefan Arold for initial OB fold structural predictions in 2012, and Dr. Sylvie Doublié for PONDR disorder predictions and discussion.

## SUPPLEMENTARY INFORMATION

**Supplementary Figure S1.**
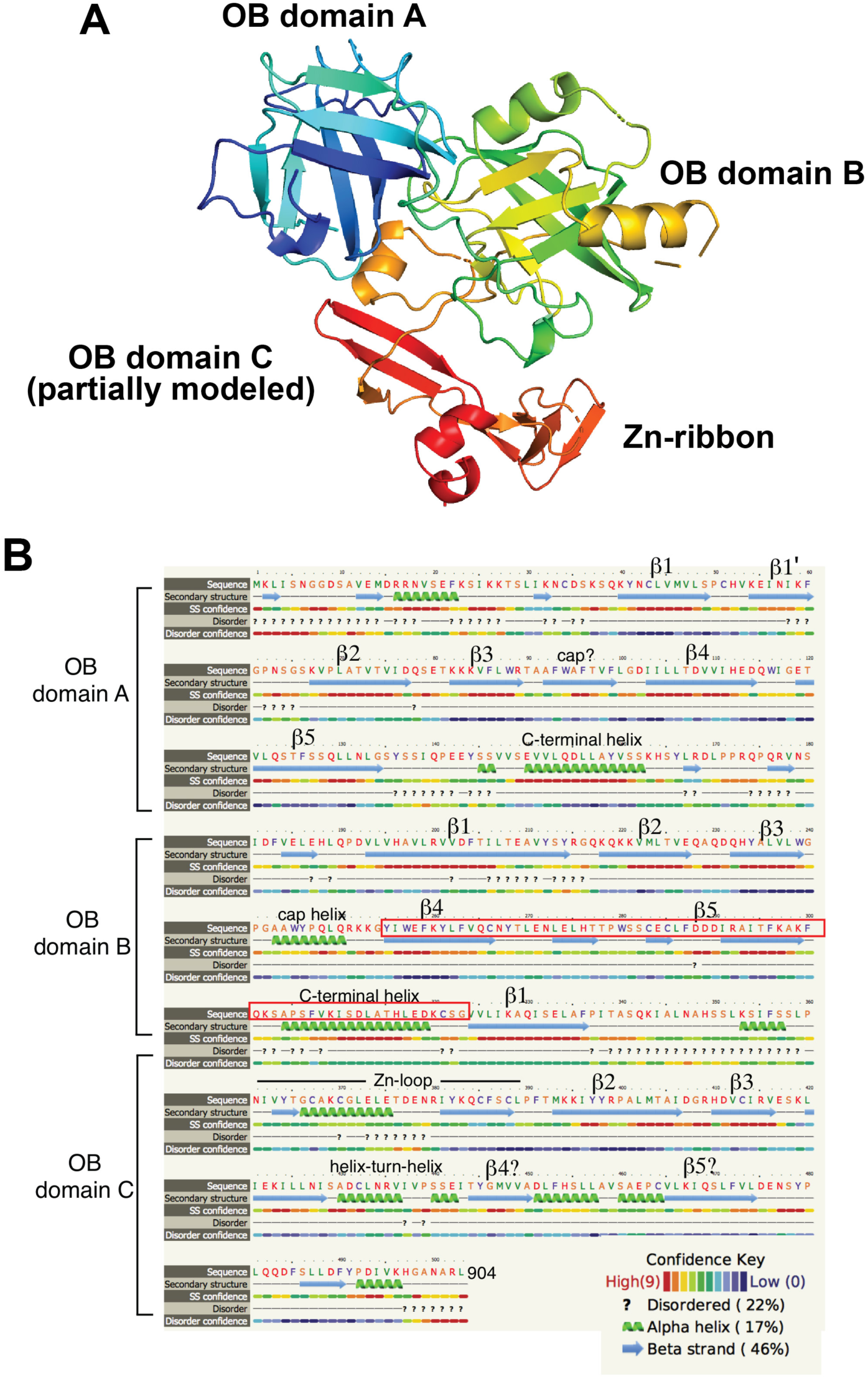
(**A**) Views of Phyre2 model for FAM35A isoform 1 residues 402-904, using template 4GOP (chain C, the RPA 70 kDa subunit from *Ustilago maydis*). The model includes OB folds with 5-stranded beta barrels for OB domain A and OB domain B, and models the first three beta strands of OB domain C, and the Zn-ribbon that occurs in a loop between OB fold C beta strands 1 and 2 (Figure S1B). (**B**) Secondary structure element assignment for the ordered region of human FAM35A (isoform 1). The secondary structure assignment is derived from the Phyre2 model (Supplementary Figure 1A) and largely coincides with features in the three functional DNA binding domains (dbdA, B and C) of RPA1 (PDB: 4GOP, 1FGU), with a few differences. In FAM35A, a C-terminal helix is predicted to be present at the end of both OB folds A and B, unlike in RPA1 where this helix only follows OB domain B. Beta strands 4 and 5 of OB domain C were not modeled by Phyre2 but candidate beta strands are present in appropriate positions, as noted. The exon absent in FAM35A isoform 2 results in 69 fewer residues (red box), deleting structural elements of OB domain B.

**Supplementary Figure S2.**
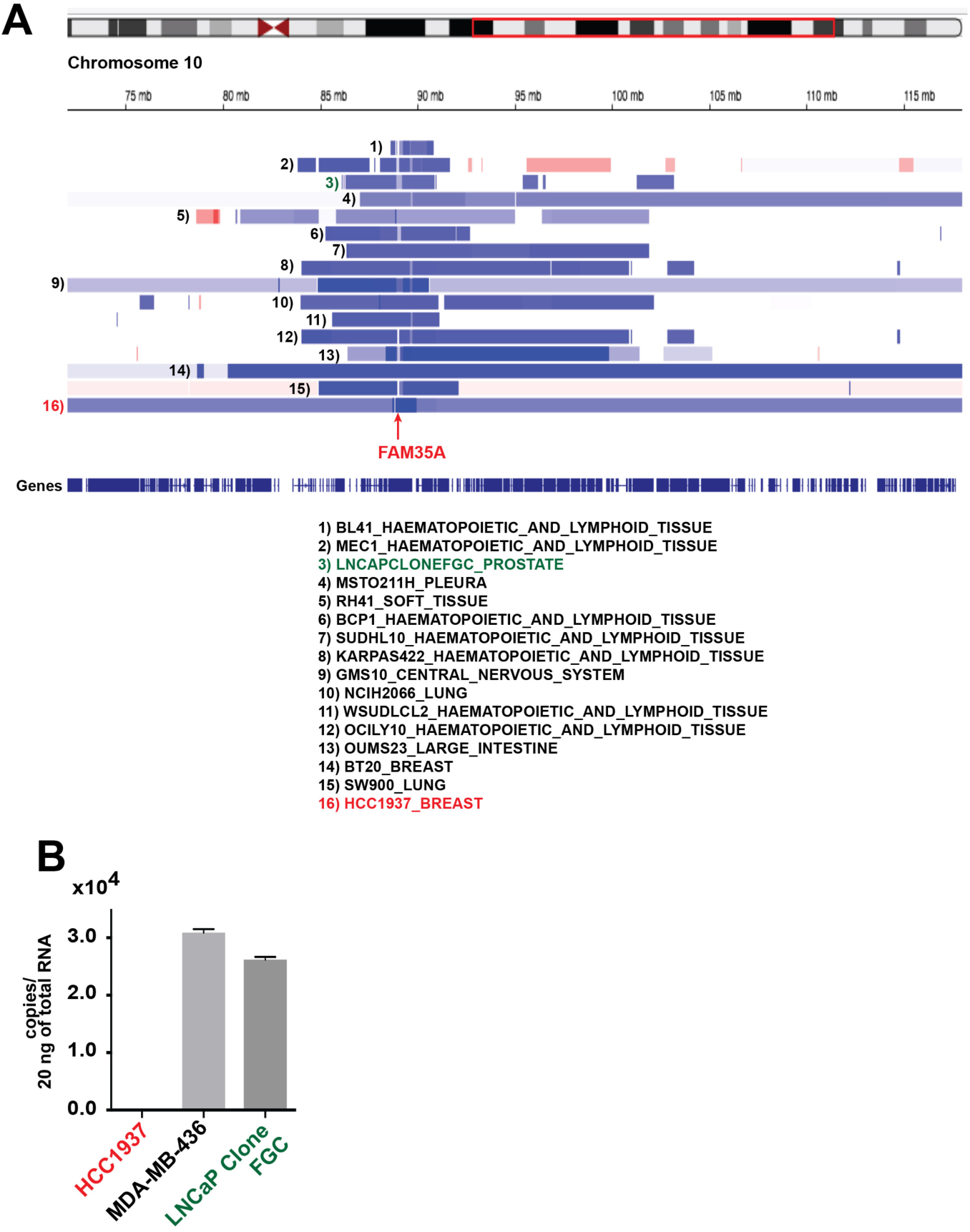
(**A**) Cell lines annotated in the Cancer Cell Line Encyclopedia with deep deletion of FAM35A. Data accessed from cBioPortal, February 2018 (**B**) Quantification of endogenous FAM35A mRNA expression. Endogenous FAM35A mRNA from HCC1937 (*BRCA1* mutant), MDA-MB-436 (*BRCA1* mutant) and LnCaP clone FGC cell lines was quantified using qPCR. Data represent mean ± SEM. n=3.

**Supplementary Figure S3.**
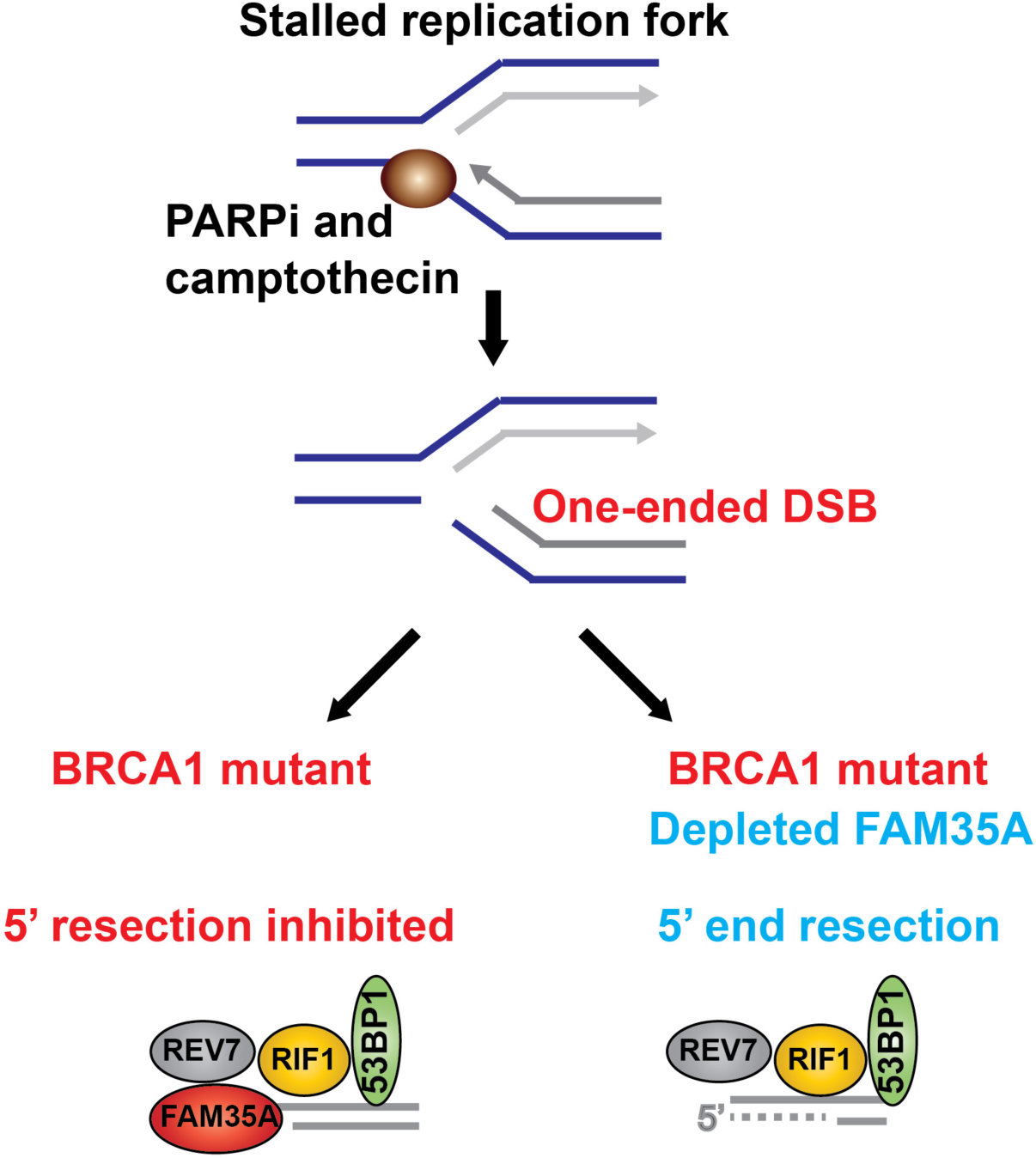
In normal cells, PARPi or camptothecin blocks the replication fork, producing a one-ended DSB. In BRCA1 mutant cells, 5’ end resection is inhibited. Finally homologous recombination pathway is activated to repair the DSB. FAM35A is involved in resection inhibition. Without FAM35A, 5’ end resection occurs in BRCA1 mutants, and then HR is activated. This may be one mechanism of PARP inhibitor chemoresistance in BRCA1 mutant cells. FAM35A may directly interact with single stranded DNA and then block nucleases or interactions with the indicated proteins.

**Supplementary Figure S4.**
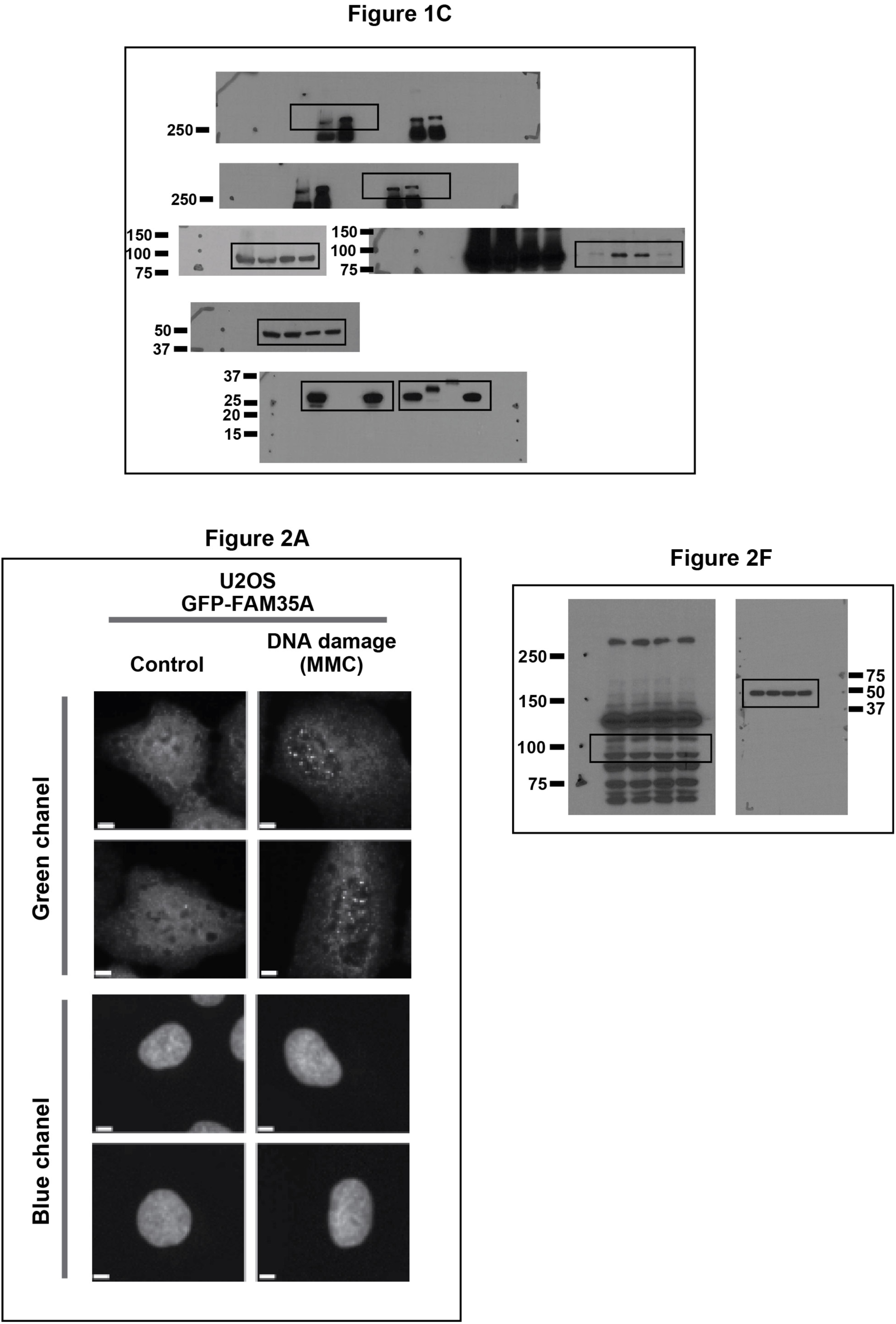

## References

1. Tomida, J. et al. REV7 is essential for DNA damage tolerance via two REV3L binding sites in mammalian DNA polymerase zeta. Nucleic Acids Res 43, 1000–11 (2015).

2. Hara, K. et al. Crystal structure of human REV7 in complex with a human REV3 fragment and structural implication of the interaction between DNA polymerase {ζ} and REV1. J Biol Chem (2010).

3. Listovsky, T. & Sale, J.E. Sequestration of CDH1 by MAD2L2 prevents premature APC/C activation prior to anaphase onset. J Cell Biol 203, 87–100 (2013).

4. Medendorp, K. et al. The mitotic arrest deficient protein MAD2B interacts with the small GTPase RAN throughout the cell cycle. PLoS One 4, e7020 (2009).

5. Hara, K. et al. Dynamic feature of mitotic arrest deficient 2-like protein 2 (MAD2L2) and structural basis for its interaction with chromosome alignment-maintaining phosphoprotein (CAMP). J Biol Chem 292, 17658–17667 (2017).

6. Pirouz, M., Pilarski, S. & Kessel, M. A critical function of Mad2l2 in primordial germ cell development of mice. PLoS Genet 9, e1003712 (2013).

7. Vermeulen, M. et al. Quantitative interaction proteomics and genome-wide profiling of epigenetic histone marks and their readers. Cell 142, 967–80 (2010).

8. Itoh, G. et al. CAMP (C13orf8, ZNF828) is a novel regulator of kinetochore-microtubule attachment. EMBO J 30, 130–44 (2011).

9. Watanabe, N. et al. The Rev7 Subunit of DNA Polymerase ζ Is Essential for Primordial Germ Cell Maintenance in the Mouse. J Biol Chem 288, 10459–71 (2013).

10. Boersma, V. et al. MAD2L2 controls DNA repair at telomeres and DNA breaks by inhibiting 5’ end resection. Nature 521, 537–40 (2015).

11. Xu, G. et al. REV7 counteracts DNA double-strand break resection and affects PARP inhibition. Nature 521, 541–4 (2015).

12. Ikura, T. et al. DNA damage-dependent acetylation and ubiquitination of H2AX enhances chromatin dynamics. Mol Cell Biol 27, 7028–40 (2007).

13. Nakatani, Y. & Ogryzko, V. Immunoaffinity purification of mammalian protein complexes. Methods Enzymol 370, 430–44 (2003).

14. Fattah, F.J. et al. The transcription factor TFII-I promotes DNA translesion synthesis and genomic stability. PLoS Genet 10, e1004419 (2014).

15. Kim, W. et al. Systematic and quantitative assessment of the ubiquitin-modified proteome. Mol Cell 44, 325–40 (2011).

16. Matsuoka, S. et al. ATM and ATR substrate analysis reveals extensive protein networks responsive to DNA damage. Science 316, 1160–6 (2007).

17. Fan, J. & Pavletich, N.P. Structure and conformational change of a replication protein A heterotrimer bound to ssDNA. Genes Dev 26, 2337–47 (2012).

18. Bochkareva, E., Korolev, S., Lees-Miller, S.P. & Bochkarev, A. Structure of the RPA trimerization core and its role in the multistep DNA-binding mechanism of RPA. EMBO J. 21, 1855–63 (2002).

19. Arunkumar, A.I., Stauffer, M.E., Bochkareva, E., Bochkarev, A. & Chazin, W.J. Independent and coordinated functions of replication protein A tandem high affinity single-stranded DNA binding domains. J Biol Chem 278, 41077–82 (2003).

20. Lin, Y.-L. et al. The evolutionarily conserved zinc finger motif in the largest sub-unit of human RPA is required for DNA replication and mismatch repair but not for nucleotide excision repair. J. Biol. Chem. 273, 1453–1461 (1998).

21. Pierce, A. et al. Comparative antiproliferative effects of iniparib and olaparib on a panel of triple-negative and non-triple-negative breast cancer cell lines. Cancer Biol Ther 14, 537–45 (2013).

22. Lehmann, B.D. et al. Identification of human triple-negative breast cancer subtypes and preclinical models for selection of targeted therapies. J Clin Invest 121, 2750–67 (2011).

23. Drew, Y. et al. Therapeutic potential of poly(ADP-ribose) polymerase inhibitor AG014699 in human cancers with mutated or methylated BRCA1 or BRCA2. J Natl Cancer Inst 103, 334–46 (2011).

24. Hill, S.J., Clark, A.P., Silver, D.P. & Livingston, D.M. BRCA1 pathway function in basal-like breast cancer cells. Mol Cell Biol 34, 3828–42 (2014).

25. Zhang, J. et al. Chk2 phosphorylation of BRCA1 regulates DNA double-strand break repair. Mol Cell Biol 24, 708–18 (2004).

26. Bunting, S.F. et al. 53BP1 inhibits homologous recombination in Brca1-deficient cells by blocking resection of DNA breaks. Cell 141, 243–54 (2010).

27. Patel, D.S., Misenko, S.M., Her, J. & Bunting, S.F. BLM helicase regulates DNA repair by counteracting RAD51 loading at DNA double-strand break sites. J Cell Biol 216, 3521–3534 (2017).

28. Densham, R.M. et al. Human BRCA1-BARD1 ubiquitin ligase activity counteracts chromatin barriers to DNA resection. Nat Struct Mol Biol 23, 647–55 (2016).

29. Mateos-Gomez, P.A. et al. Mammalian polymerase θ promotes alternative NHEJ and suppresses recombination. Nature 518, 254–7 (2015).

30. Wyatt, D.W. et al. Essential Roles for Polymerase theta-Mediated End Joining in the Repair of Chromosome Breaks. Molecular Cell 63, 662–73 (2016).

31. Chaudhuri, A.R. et al. Replication fork stability confers chemoresistance in BRCA-deficient cells. Nature 535, 382–7 (2016).

32. Romano, P. et al. Cell Line Data Base: structure and recent improvements towards molecular authentication of human cell lines. Nucleic acids research 37, D925–32 (2009).

33. Tomida, J. et al. A novel interplay between the Fanconi anemia core complex and ATR-ATRIP kinase during DNA cross-link repair. Nucleic Acids Res 41, 6930–41 (2013).

34. Takata, K., Reh, S., Tomida, J., Person, M.D. & Wood, R.D. Human DNA helicase HELQ participates in DNA interstrand crosslink tolerance with ATR and RAD51 paralogs. Nature Communications 4(2013).

35. Galanty, Y., Belotserkovskaya, R., Coates, J. & Jackson, S.P. RNF4, a SUMO-targeted ubiquitin E3 ligase, promotes DNA double-strand break repair. Genes Dev 26, 1179–95 (2012).

36. Lou, Z. et al. MDC1 regulates DNA-PK autophosphorylation in response to DNA damage. J Biol Chem 279, 46359–62 (2004).

37. Lee, Y.S., Gregory, M.T. & Yang, W. Human Pol zeta purified with accessory subunits is active in translesion DNA synthesis and complements Pol eta in cisplatin bypass. Proc Natl Acad Sci U S A 111, 2954–9 (2014).

38. Takata, K., Reh, S., Tomida, J., Person, M.D. & Wood, R.D. Human DNA helicase HELQ participates in DNA interstrand crosslink tolerance with ATR and RAD51 paralogs. Nat Commun 4, 2338 (2013).

39. Nakatani, Y. & Ogryzko, V. Immunoaffinity purification of mammalian protein complexes. Methods in enzymology 370, 430–44 (2003).

40. Fradet-Turcotte, A. et al. 53BP1 is a reader of the DNA-damage-induced H2A Lys 15 ubiquitin mark. Nature 499, 50–4 (2013).

41. Escribano-Diaz, C. et al. A cell cycle-dependent regulatory circuit composed of 53BP1-RIF1 and BRCA1-CtIP controls DNA repair pathway choice. Mol Cell 49, 872–83 (2013).

42. Kelley, L.A., Mezulis, S., Yates, C.M., Wass, M.N. & Sternberg, M.J.E. The Phyre2 web portal for protein modeling, prediction and analysis. Nature Protocols 10, 845 (2015).

43. Gao, J. et al. Integrative analysis of complex cancer genomics and clinical profiles using the cBioPortal. Sci Signal 6, pl1 (2013).

44. Rhodes, D.R. et al. Oncomine 3.0: genes, pathways, and networks in a collection of 18,000 cancer gene expression profiles. Neoplasia 9, 166–80 (2007).

45. Warnat, P. et al. Cross-study analysis of gene expression data for intermediate neuroblastoma identifies two biological subtypes. BMC Cancer 7, 89 (2007).

46. Grasso, C.S. et al. The mutational landscape of lethal castration-resistant prostate cancer. Nature 487, 239–43 (2012).

47. Lapointe, J. et al. Gene expression profiling identifies clinically relevant subtypes of prostate cancer. Proc Natl Acad Sci U S A 101, 811–6 (2004).

48. Taylor, B.S. et al. Integrative genomic profiling of human prostate cancer. Cancer Cell 18, 11–22 (2010).

49. Vanaja, D.K., Cheville, J.C., Iturria, S.J. & Young, C.Y. Transcriptional silencing of zinc finger protein 185 identified by expression profiling is associated with prostate cancer progression. Cancer Res 63, 3877–82 (2003).

50. Varambally, S. et al. Integrative genomic and proteomic analysis of prostate cancer reveals signatures of metastatic progression. Cancer Cell 8, 393–406 (2005).

